# A Novel Regulation on the Developmental Checkpoint Protein Sda that Controls Sporulation and Biofilm Formation in *Bacillus subtilis*

**DOI:** 10.1101/2024.05.13.593929

**Authors:** Yinghao He, Yuxuan Qin, Jennifer Greenwich, Samantha Balaban, Migue Van Louis Darcera, Kevin Gozzi, Yunrong Chai

## Abstract

Biofilm formation by *Bacillus subtilis* is triggered by an unusually simple environmental sensing mechanism. Certain serine codons, the four TCN codons (N for A, T, C, or G), in the gene for the biofilm repressor SinR caused lowered SinR translation and subsequent biofilm induction during transition from exponential to stationary growth. Global ribosome profiling showed that ribosomes pause when translating the four UCN (U for T on the mRNA) serine codons on mRNA, but not the two AGC/AGU serine codons. We proposed a serine codon hierarchy (AGC/AGT vs TCN) in that genes enriched in the TCN serine codons may experience reduced translation efficiency when serine is limited. In this study, we designed an algorithm to score all protein-coding genes in *B. subtilis* NCIB3610 based on the serine codon hierarchy. We generated a short list of 50 genes that could be subject to regulation by this novel mechanism. We further investigated one such gene from the list, *sda*, which encodes a developmental checkpoint protein regulating both sporulation and biofilm formation. We showed that synonymously switching the TCN serine codons to AGC in *sda* led to delayed biofilm formation and sporulation. This engineered strain also outgrew strains with other synonymously substituted *sda* alleles (TCN) in competition assays for biofilm formation and sporulation. Lastly, we showed that the AGC serine codon substitutions in *sda* elevated the Sda protein levels. This serine codon hierarchy-based novel signaling mechanism could be exploited by bacteria in adapting to stationary phase and regulating important biological processes.

**Importance:** Genome-wide ribosome profiling in *Bacillus subtilis* shows that under serine limitation, ribosomes pause on the four TCN (N for A, C, G, and T), but not AGC/AGT serine codons, during translation at a global scale. This serine codon hierarchy (AGC/T vs TCN) differentially influences translation efficiency of genes enriched in certain serine codons. In this study, we designed an algorism to score all 4000+ genes in the *B. subtilis* genome and generated a list of 50 genes that could be subject to this novel serine codon hierarchy-mediated regulation. We further investigated one such gene, *sda*, encoding a developmental check point protein. We show that *sda* and cell developments controlled by Sda are also regulated by this novel mechanism.

## Introduction

Biofilm formation is a developmental process in which planktonic cells switch from vegetative growth to assembly of multicellular communities in response to environmental cues (Flemming *et al*., 2016, O’Toole *et al*., 2000, López *et al*., 2010, Sauer *et al*., 2022). Within the biofilm, cells are encased by a self-produced, extracellular matrix to protect them from toxins and chemicals with adverse effects (Flemming & Wingender, 2010, Arnaouteli *et al*., 2021). Cells in the biofilm are also highly differentiated, displaying phenotypically distinct subpopulations (Lopez *et al*., 2009, van Gestel *et al*., 2015, Qin *et al*., 2022). In the model bacterium *Bacillus subtilis*, biofilm formation is initiated when the master regulator Spo0A is activated by protein phosphorylation (Spo0A∼P) via a multi-component phosphor-relay, controlled by a crew of sensory histidine kinases (from KinA to KinE, Figure 1)(Vlamakis *et al*., 2013, Cairns *et al*., 2014, Hoch, 1993). These kinases sense environmental and physiological signals and activate Spo0A via the phosphor-relay. Spo0A∼P then turns on the gene for SinI, an antagonist of the biofilm repressor SinR, and allows expression of SinR-repressed genes, which ultimately leads to biofilm induction (Chai *et al*., 2008, Kearns *et al*., 2005, Bai *et al*., 1993, Milton *et al*., 2020).

**Figure 1.**
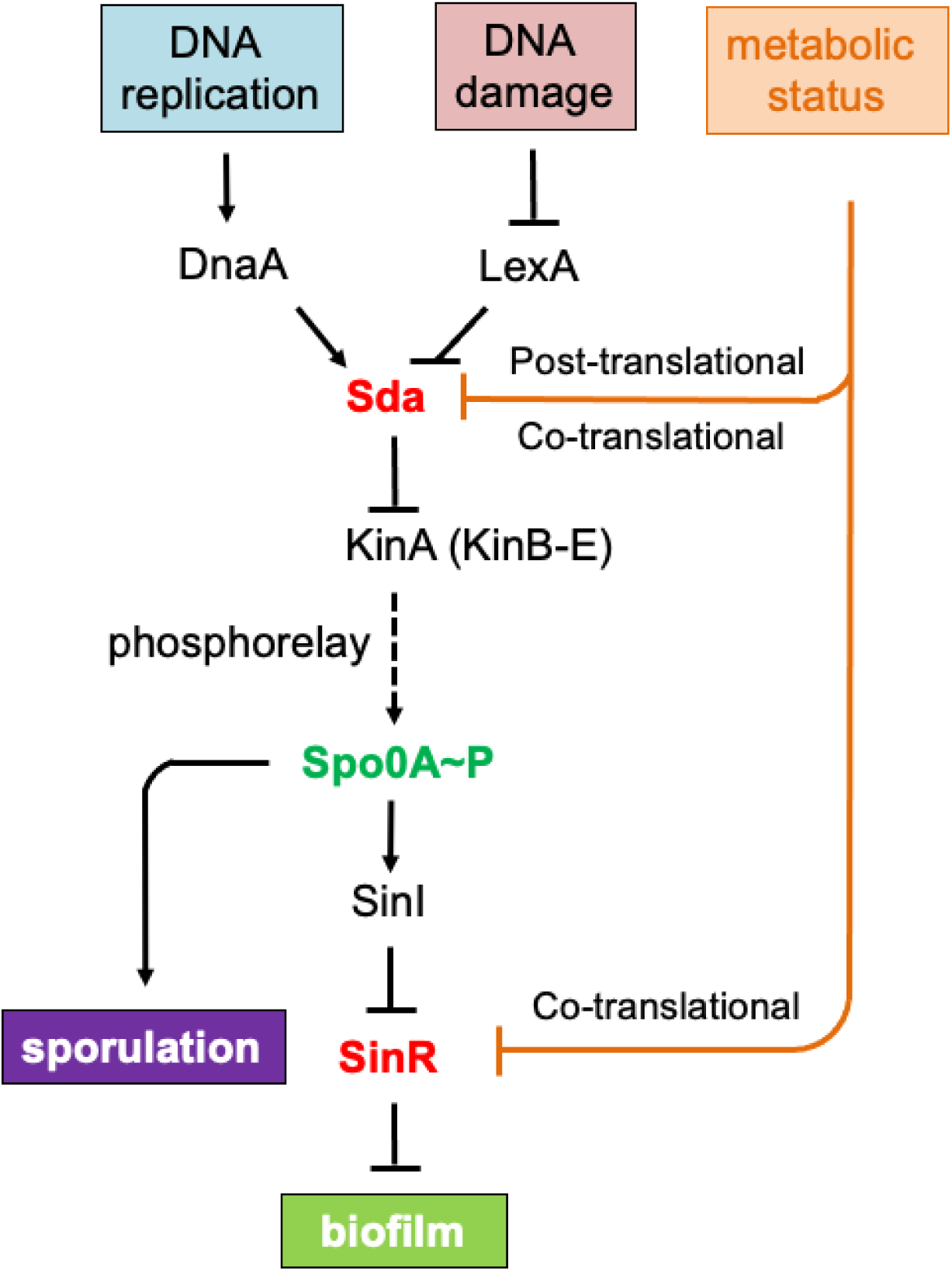
Regulation of biofilm formation and sporulation in *B. subtilis*. Biofilm formation and sporulation are both controlled by the master regulator Spo0A. Spo0A is activated by protein phosphorylation. Once phosphorylated, Spo0A∼P turns on *sinI*, whose protein product antagonizes the biofilm master repressor, SinR, derepresses SinR-controlled biofilm genes to initiate biofilm formation. When accumulated at high levels, Spo0A∼P also turns on hundreds of genes dedicated to spore formation. The phosphorylation of Spo0A is mediated by a crew of sensory histidine kinases (from KinA to KinE) that phosphorylate Spo0A directly or indirectly through the phosphorelay in response to environmental cues or physiological signals. A checkpoint protein, Sda, whose expression is monitored by cell growth status (through DnaA) or DNA damage (through LexA), negatively regulates Spo0A phosphorylation by blocking the transfer of the phosphoryl group from the histidine kinase to Spo0A. Here we propose a novel, serine codon hierarchy-based regulatory mechanism, previously shown to regulate the *sinR* gene, also regulates *sda* concertedly in triggering biofilm formation and sporulation in *B. subtilis*.

Phosphorylation of Spo0A is also controlled by a developmental checkpoint protein, Sda (Burkholder *et al*., 2001, Veening *et al*., 2009). Sda blocks the histidine kinase, KinA, from transferring the phosphoryl group to the phosphor-relay proteins, which activates Spo0A (Figure 1)(Rowland *et al*., 2004, Whitten *et al*., 2007). Deletion of the *sda* gene triggers sporulation much earlier than normally seen due to premature activation of Spo0A (Burkholder *et al*., 2001). Sda is regulated both transcriptionally and post-translationally (Veening *et al*., 2009, Ruvolo *et al*., 2006). At the transcriptional level, the *sda* gene is controlled by the DNA replication initiation protein DnaA in response to cell growth status; in fast growing cells, DnaA directly binds to the promoter of *sda* and activates its expression (Veening *et al*., 2009). Sda in turn blocks activation of Spo0A and sporulation so that cells are kept in vegetative growth. The *sda* gene is also a member of the DNA damage response (DDR) regulon in *B. subtilis*, and is negatively regulated by LexA, a master repressor for DDR (Au *et al*., 2005, Gozzi *et al*., 2017). Regulation by LexA ensures that if there is elevated DNA damage in the cell, Sda protein is produced, blocking Spo0A activation and initiation of sporulation. Such a checkpoint mechanism prevents the packaging of damaged DNA into a forespore, allowing time for repairing detrimental mutations. In addition to transcriptional regulations, Sda protein levels decline when cells enter the stationary phase due to proteolysis (Ruvolo *et al*., 2006). This further diminishes Sda activity and triggers Spo0A activation as discussed above.

Constant monitoring of changing environments and internal physiological states is essential to bacterial survival. Studies have revealed various sensing mechanisms in bacteria, involving either dedicated protein sensors or riboswitch-like RNAs (West & Stock, 2001, Henkin & Yanofsky, 2002). For example, when initiating biofilm formation in *B. subtilis*, multiple histidine kinases serve as receptors to transduce signals from environments (Vlamakis *et al*., 2013, Chen *et al*., 2012, McLoon *et al*., 2011, López *et al*., 2009, Shemesh & Chai, 2013, Kolodkin-Gal *et al*., 2013, Beauregard *et al*., 2013). RNA-based riboswitches are also well-known sensors for nutrient molecules such as amino acids and vitamins, and they often regulate the activity of the respective biosynthetic genes (Henkin & Yanofsky, 2002, Breaker, 2012). In a previous study (Subramaniam *et al*., 2013), we revealed a novel signaling mechanism for biofilm induction in *B. subtilis* that relies on neither dedicated protein sensors nor RNAs, but rather a serine codon hierarchy identified through studying global translation dynamics. Specifically, the uncommon ratio and distribution of certain synonymous serine codons in the gene encoding the biofilm master repressor SinR were found to impact the translation efficiency of *sinR* under serine starvation.

In bacteria, there are six synonymous triplet codons for the amino acid serine: AGC, AGT, TCA, TCC, TCG, and TCT. In our previous study using global ribosome profiling (Subramaniam *et al*., 2013), we observed that when cells enter stationary phase or during biofilm induction (conditions that coincide with serine depletion), ribosomes pause when translating the UCN (N for A, C, G, or U) serine codons but not when translating the AGC/AGU serine codons. The observed ribosome pause on the UCN serine codons is a global phenomenon. However, it happens only during serine starvation (e.g. in stationary phase or biofilm induction), but not in log phase or when excess serine is supplemented to the media (Subramaniam *et al*., 2013). These results suggest that the six synonymous serine codons can be divided into two different groups based on their influence on global protein translation (AGC/T as “good” vs TCA/C/G/T as “bad”). Subtle differences in codons within each group allowed us to further propose a serine codon hierarchy (AGC>AGT>TCA>TCT=TCC=TCG).

Interestingly, the hierarchy of synonymous serine codons does not correlate with genome-wide serine codon usage in *B. subtilis* (AGC:22.5%, AGT:10.6%, TCA: 23.7%, TCC:12.8%, TCG: 10.2%, and TCT: 20.3%)(Moszer *et al*., 1999). Biased codon usage is a universal phenomenon across different domains of life. It has been proposed that the ratio of rare codons (codons with a much lower genome-wide usage frequency) in a gene could impact the translation efficiency of the gene (Liu *et al*., 2021). Surprisingly, in our global ribosome profiling, we observed little or no impact on the ribosome translation speed that can be correlated to the statistically determined codon usage for any amino acid. On the other hand, the serine codon hierarchy that we characterized clearly causes ribosome pause and is nutrient-status dependent; the four TCN serine codons slow down ribosome movement when serine is exhausted (Subramaniam *et al*., 2013).

How does this serine codon hierarchy specifically influence biofilm induction in *B. subtilis*? Interestingly, the *sinR* gene encoding the biofilm repressor contains twice the number of the overall serine codons than the genome average as well as a strong bias toward the TCN serine codons (Pedreira *et al*., 2022). This puts *sinR* one of the potential candidates whose translation could be strongly influenced during serine starvation; ribosomes pause more frequently when translating those UCN codons on the *sinR* mRNA, which results in lowered SinR protein levels and ultimately derepression of biofilm genes controlled by SinR (Subramaniam *et al*., 2013). In a follow-up study, we presented evidence on why the amino acid serine is depleted during the transition from log to stationary phase (which again coincides with biofilm induction), and why ribosomes pause more frequently on the UCN serine codons than the AGC/AGU codons when supply of serine becomes limited (Greenwich *et al*., 2019).

Although we expect this serine codon hierarchy-based signaling mechanism to be broadly applicable, so far it has only been shown to regulate *sinR* and influence SinR-controlled biofilm induction in *B. subtilis* (Subramaniam et al., 2013). Whether it impacts translation efficiency of other genes (with a similar ratio and distribution of synonymous serine codons) and other biological processes in *B. subtilis* remains unknown. In this study, we first designed an algorithm that allowed us to score all protein-coding genes in the *B. subtilis* NCIB3610 genome based on the ratio and distribution of the 6 synonymous serine codons (see Methods). We identified a list of 50 genes that bear a much higher ratio of serine codons than the genome average and a strong bias toward the TCN codons. We then picked one such gene, *sda*, encoding a developmental checkpoint protein. We showed that similar to *sinR*, the serine codon hierarchy also influences the activity of *sda* and Sda-controlled sporulation and biofilm development in *B. subtilis*.

## Results

A bioinformatics search reveals genes in the *B. subtilis* genome potentially influenced by the serine codon hierarchy.

We speculated that in addition to *sinR*, the serine codon hierarchy could influence translation efficiency of other genes as well under serine starvation. We designed a scoring system based on the ratio of overall serine codons and the bias toward specific serine codons (TCN or AGC/T), which allowed us to score each of the >4000 protein-encoding genes in the genome of *B. subtilis* NCIB3610 (Nye *et al*., 2017).

Each of the six serine codons was given a score based on previously observed ribosome occupancy on each serine codon across the genome during serine starvation (Subramaniam *et al*., 2013). We came up with a list of 50 genes, all of which with a much higher than average ratio of serine codons and a strong bias toward the TCN codons (Table S1). The percentages of serine codons in these 50 genes vary from the highest of 24.43% and the lowest of 9.53%, compared to the genome average of 6.3% in *B. subtilis (Subramaniam et al., 2013)*. We also categorized these genes based on their known or predicted functions (Table 1). Among these 50 genes, 15 are sporulation-related genes. There are also a significant number of genes (15) predicted to encode either secreted or membrane proteins. In addition, there are genes apparently in the same operons (e.g. *ydgA-ydgB, gerPB*-*PD-PE-PF*). The *sinR* gene that we characterized in the previous study is ranked No. 45 in the list. Another gene with a known regulatory function, *sda,* is ranked No. 19. The *sda* gene has a much higher percentage of serine codons than the genome average (13.2% vs 6.3%). Moreover, among the 7 serine codons in the gene, there is only one AGC codon while the other 6 are all TCN serine codons (Figure 2A).

**Figure 2.**
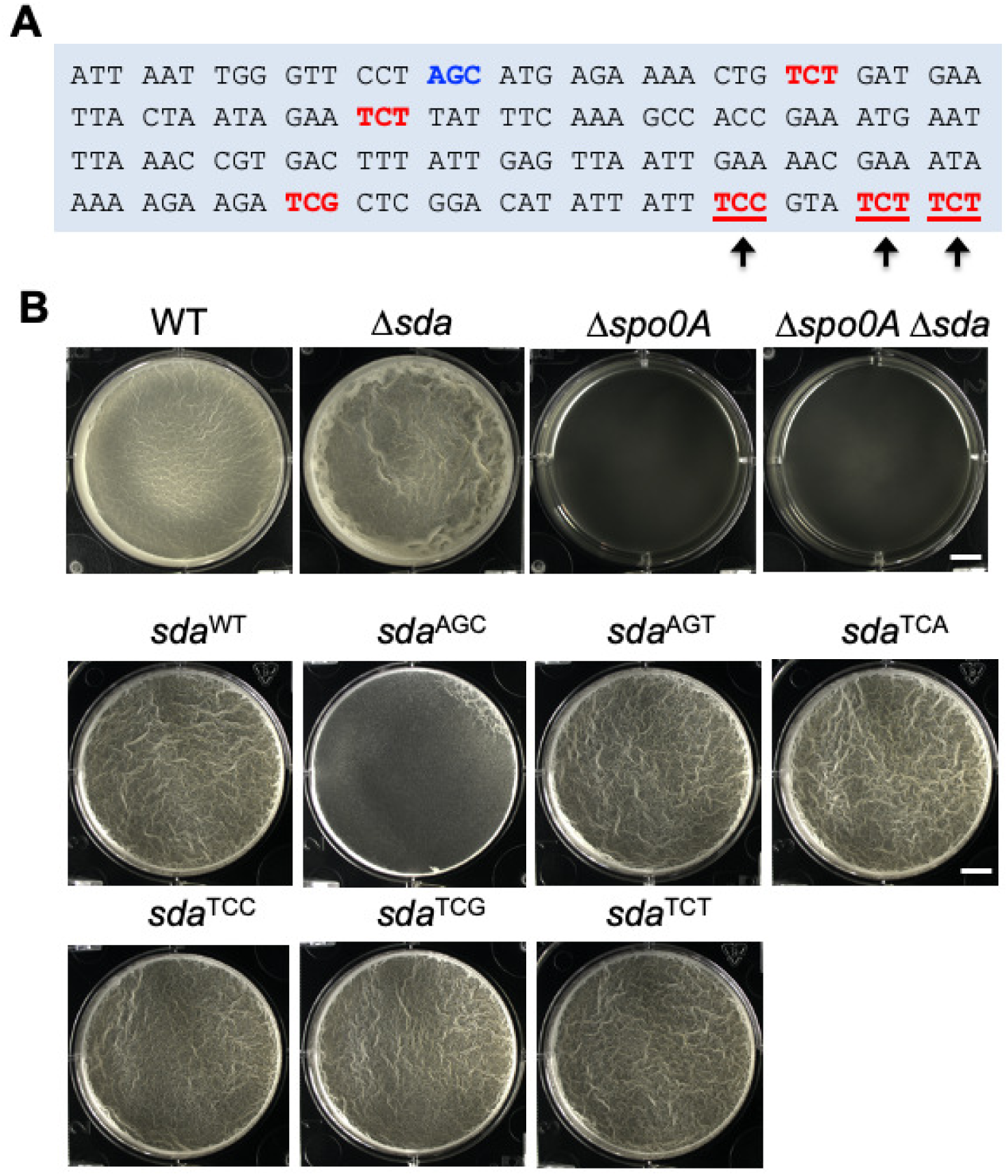
Switching synonymous serine codons in *sda* influences biofilm formation. **(A)** Shown is the *sda* coding sequence, in which the 7 serine codons are highlighted either in red (TCN codons) or blue (AGC/AGT). Six mutant *sda* alleles were constructed, each with one of the six synonymous substitutions for the terminal three serine codons (indicated by arrows). **(B)** Formation of pellicle biofilms in MSgg by the wild type, *Δsda*, *Δspo0A*, *Δspo0A Δsda*, and 7 complemented strains with the deletion in the native *sda* gene and complementation at *amyE* by either one of the six synonymously substituted *sda* alleles or the wild-type *sda*. Cells were incubated at 30°C for 48 hours before images were taken. Scale bars, 250 μm. Scale bars are representative to all panels in Figure 2B.

**Table 1.**
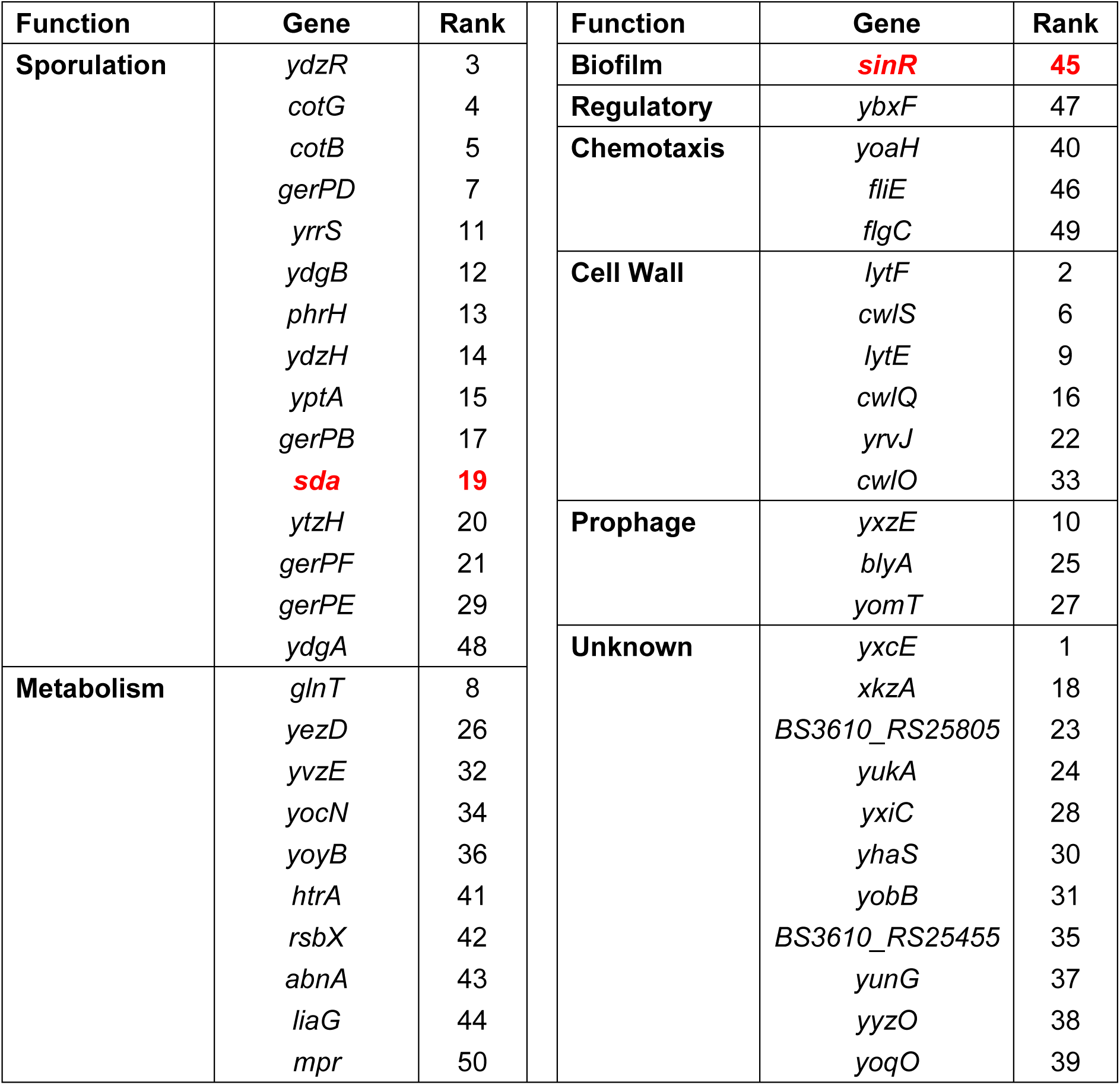
The top 50 genes with biased serine codon usage in *B. subtilis*.

### Switching synonymous serine codons in *sda* impacts biofilm formation

Sda is a known developmental checkpoint protein that controls initiation of sporulation by blocking premature activation of Spo0A (Burkholder *et al*., 2001, Veening *et al*., 2009). Sda is also proposed to regulate biofilm formation since its target, Spo0A, governs both sporulation and biofilm formation in *B. subtilis* (Gozzi et al., 2017). An *sda* insertional deletion mutant (Δ*sda*) was constructed. This mutant formed an early and more robust pellicle biofilm with visible wrinkles than the wild type on day two (Figure 2B). We also confirmed that the effect of Sda on biofilm robustness is mediated by Spo0A since a double mutant of Δ*spo0A*Δ*sda* is phenotypically identical to the *spo0A* single mutant (Figure 2B).

Since both *sinR* and *sda* are among the top 50 genes in our research, we predicted that like *sinR*, ribosomes may pause more frequently when translating the UCN serine codons on the *sda* mRNA during serine starvation. Reduced translation efficiency, in concerted action with other known transcriptional or post-translational regulations on *sda* reported in previous studies (Ruvolo *et al*., 2006, Veening *et al*., 2009, Gozzi *et al*., 2017), may quickly diminish the activities of Sda, which ultimately leads to Spo0A activation as we discussed above. In previous studies, we defined AGC/T as “good” serine codons whereas TCN as “bad” based on their influence on translational efficiency during serine starvation (Subramaniam *et al*., 2013, Greenwich *et al*., 2019). When systematically switching the AGC/T codons in *sinR* to TCN codons or *vice versa*, we observed detectable alterations in SinR protein abundance and the biofilm phenotypes, suggesting that synonymous changes in the serine codons in *sinR* are not neutral in SinR activities (Subramaniam *et al*., 2013).

We decided to similarly examine the impact of synonymous serine codon changes in *sda*. To do so, three terminal TCN serine codons in *sda* (5’-<u>TCC</u> GTA <u>TCT</u> <u>TCT</u>-3’, Figure 2A) were substituted with each of the six synonymous serine codons. Each substituted allele of *sda* was introduced into the *amyE* locus of the Δ*sda* mutant. We then tested the biofilm phenotypes of the engineered strains. We expect that the strains with the AGC/T codon substitutions may synthesize more Sda protein under biofilm-inducing conditions (serine limitation) and thus form weaker biofilms, while strains with the TCN codon substitutions likely produce similar amounts of Sda seen in the wild type. As shown in Figure 2B (the second and third rows), biofilms formed by the 4 strains with the TCN substitutions (TCA, TCC, TCG, and TCT) were indistinguishable to that of the wild type (*sda*^WT^). On the other hand, the strain with the AGC substitution (but surprisingly not the AGT substitution) formed a weaker biofilm. According to the proposed serine codon hierarchy (AGC>AGT>TCA≥TCT=TCC=TCG), the AGC codon has the highest translation efficiency during serine starvation (Subramaniam *et al*., 2013). Our results are consistent with that prediction even though the AGT substitution showed little difference form the wild type (Figure 2B).

### Switching synonymous serine codons in *sda* impacts the timing of sporulation

We wondered whether switching synonymous serine codons in *sda* also impacts sporulation. Under laboratory settings, *B. subtilis* sporulation is an 8-hour process (past *T*_0_) that can be divided into several morphologically distinct stages (Figure 3A)(Tan & Ramamurthi, 2014, Errington, 1993). This asymmetric cell division process, triggered by activation of Spo0A, produces a small cell (forespore) and a large cell (mother cell). The asymmetric septum then curves out to initiate engulfment from the large cell to the forespore. By staining cells with a membrane dye (FM4-64), we visualized vegetative growth, asymmetric division, forespore, engulfment, and lastly the endospore (Figure 3A). This also allowed us to quantify individual cells in each sporulation stage. We picked three different strains, the wild type, the strain with the AGC serine codon substitutions in *sda*, and the strain with the TCC serine codon substitutions, and compared synchronized sporulating cell populations by the three strains. We found that the timing that the morphologically distinct stages appeared during sporulation differed significantly among the three strains. For example, after six hours of growth from inoculation (around *T*_1_), 62.9% of the cells of the *sda*^AGC^ strain were still in vegetative growth, a ratio almost three times higher than that in the wild type (20.7%) and the *sda*^TCC^ strain (26.4%), indicating a delay in sporulation in the *sda*^AGC^ strain (Figure 3B-C). Further, the engulfment of the forespore was rarely seen in the *sda*^AGC^ strain, but frequently observed in both the wild type and the *sda*^TCC^ strain (Figure 3B-C). Based on these observations, we speculated that under the same synchronized sporulating conditions, the activities of Sda were diminished at a slower pace in the *sda*^AGC^ strain than in the wild type and the *sda*^TCC^ strain, causing a delay in Spo0A activation and initiation of sporulation in the *sda*^AGC^ strain. This delay was also seen when cells were collected from another time point (Figure S1).

**Figure 3.**
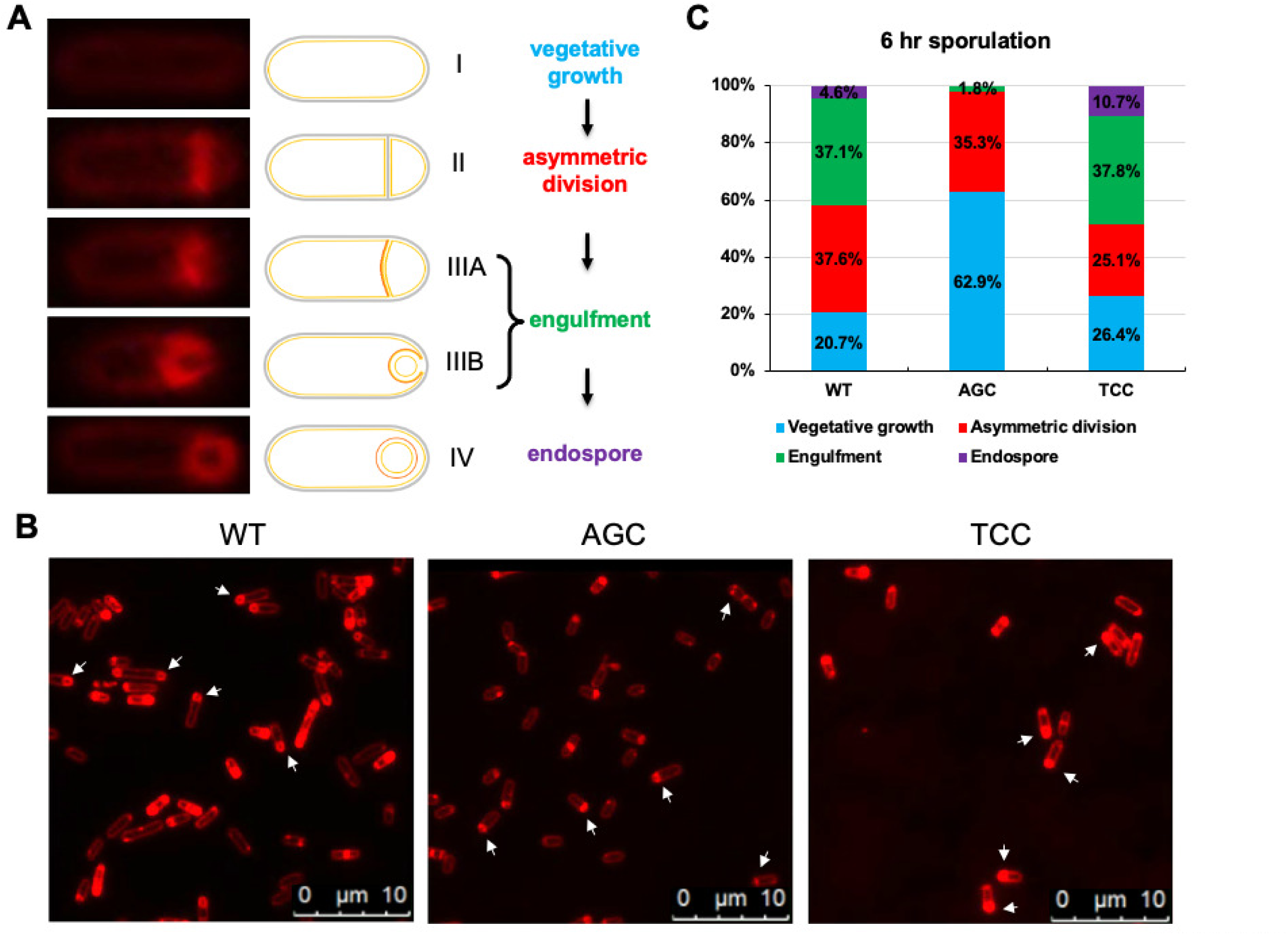
Switching synonymous serine codons in *sda* alters the timing of sporulation. **(A)** Representative *B. subtilis* cells in different developmental stages (vegetative growth, asymmetric division, engulfment, and endospore) during sporulation are included, accompanied by a diagram. Cells were stained with the membrane dye FM4-64. From stage I to II, cells shift from vegetative growth to the formation of asymmetric division septum at one polar. In stages II and III (A and B), the asymmetric division results in two compartments: forespore and mother cell. In stage IV, the mother cell engulfs the forespore. **(B)** Stain of the cell membrane by FM4-64. Cells were grown with shaking at 37°C for 6 hours in DMS before harvested and stained with the dye. Strains used here include the wild type (WT), and Δ*sda* complemented at *amyE* with either *sda*^AGC^ (AGC) or *sda*^TCC^ (TCC). Arrows point to the cells with the highest ratio at a specific stage in each panel. Scale bars shown are 10 μm. **(C)** Quantification of the ratio of cells in each of in four sporulation stages: I, vegetative growth (blue); II, asymmetric division (red); III, engulfment (green); IV endospore (purple). Approximately 300 cells were randomly picked and analyzed for each sample.

Next, we carried out heat kill experiments to further test whether the observed delay in sporulation caused by switching synonymous serine codons in *sda* could impact the timing of forming heat-resistant spores. We harvested cells of the wild type, the *sda*^AGC^ strain, and the *sda*^TCC^ strain at different times post the initiation of sporulation (*T*_0_), and performed heat kill assays. Our results (Figure 4A) show that at *T*_10_ (10 hours after sporulation initiation), the wild-type culture contained a significant amount of heat-resistant spores (58%) while the *sda*^AGC^ culture had a very low ratio of heat-resistant spores (5%). The ratio of heat-resistant spores in the *sda*^TCC^ culture was slightly lower than that of the wild type (44% vs 58%, Figure 4A). Similar differences were also seen in cells harvested at *T*_8_ albeit ratios of heat-resistant spores were dramatically lower in all three strains (Figure 4B). The above results strongly suggest that the impact of Sda activities on the timing of sporulation initiation could be critically important under certain circumstances. The foreseeable benefit of having many “bad” TCN codons in a gene coding for a developmental checkpoint protein like Sda is to ensure timely initiation of sporulation for cells to survive adverse environmental conditions.

**Figure 4:**
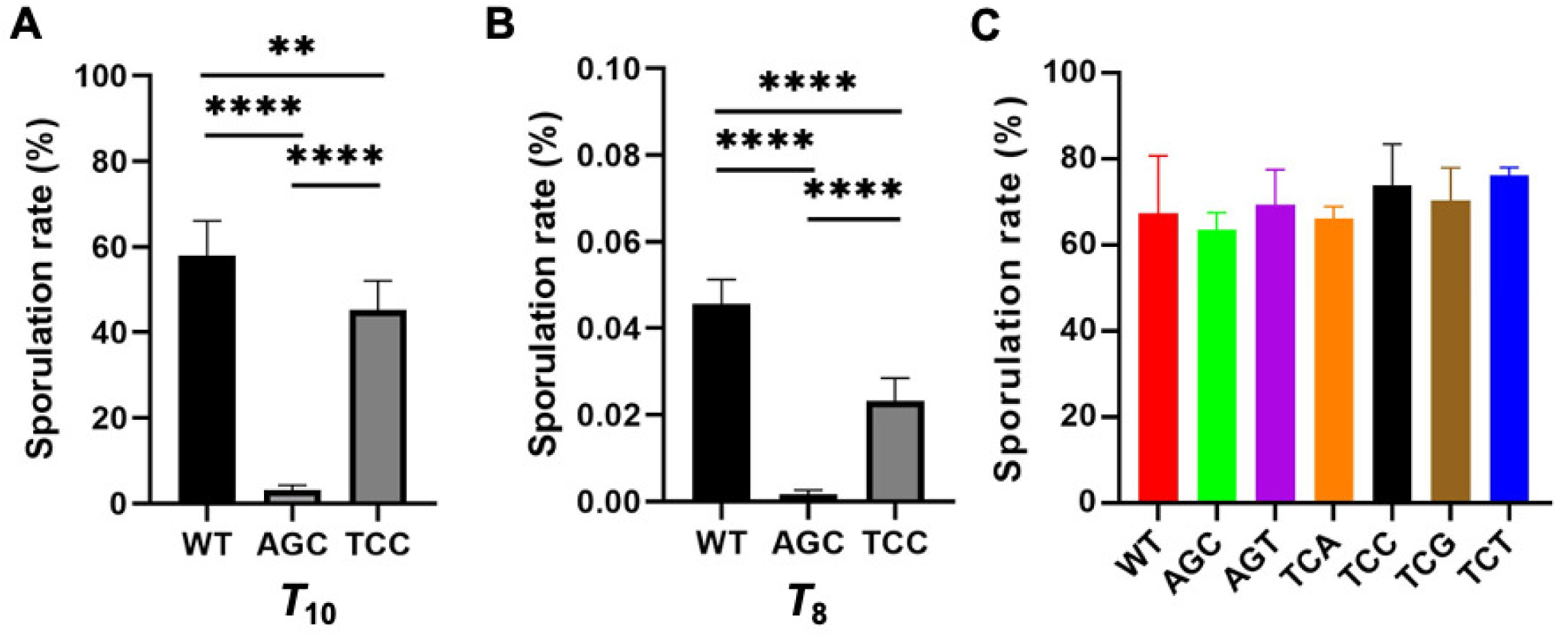
The AGC mutant had a delayed sporulation compared to the wild-type and the TCC mutant. **(A-B)** The sporulation efficiency of the wild-type and the AGC and TCC mutants was compared by quantifying sporulation rate of each strain at 6, 8, and 10 hours (post *T_0_*), respectively. Shown here are sporulation results of 8 (A) and 10 hr (B) samples (6 hr samples had very low sporulation rate and results are not shown here). Fresh cultures of each strain were normalized to OD_600_∼1, then 1 to 100 inoculated into DSM medium, and grown in shaking at 37°C. At each time point, CFUs of each strain were quantified before and after heat-kill. Sporulation rate was calculated by dividing CFU of pre-heat kill culture by CFU of after heat-kill culture. **** indicates *p* value <0.0001, and ** indicates *p* value <0.01. Statistical analysis was performed by the unpaired t-test from Prism. (C) Terminal sporulation rate between the wild-type strain and all six synonymous serine codon mutants showed insignificant differences. Cells of each strain were grown to mid log phase and inoculated 1 to 100 into DSM medium. After incubation at 37°C for 30 hours, cells were collected and heat kill assay was performed. Sporulation rate was calculated similarly as described above. The differences among different strains are insignificant.

All synonymous codon switching strains had a similar terminal sporulation efficiency to that of the wild type after grown in sporulation media for 30 hours (Figure 4C), reinforcing the idea that the codon switch caused a delay in the timing of sporulation, not a damage to sporulation capacity. Lastly, we tested the growth of all engineered strains in both the sporulation (DSM) and the LB growth media, no difference was observed (Figure S2). This ruled out the possibility that delay in sporulation in the engineered strain is due to a delay in growth. The above results again support the idea that synonymously switching serine codons in *sda* alters the activities of Sda and thus the timing of sporulation.

### Sporulation and biofilm competition assays among the *sda* alleles with synonymous serine codon changes

The phenotypical changes in both biofilm formation and sporulation caused by the synonymous serine codon changes in *sda* were modest (Figures 2-4). We thus designed two competition assays, one for biofilm formation and the other for sporulation, to further examine the differences caused by different *sda* alleles. We reasoned that the co-cultured synonymous mutants of *sda* would compete and reveal subtle differences in fitness. Competition assays would allow us to identify the mutants with certain synonymous substitutions in *sda* that outcompete/outgrow other strains in the co-cultured biofilm community or during sporulation. The relative proportion of particular mutant(s) will increase and become dominant after several rounds of competition, and *vice versa* for the ones that lose competitiveness.

We inoculated equal numbers of cells from the six serine synonymous mutants of *sda* as well as the wild-type strain into the biofilm (MSgg) and the sporulation media (DSM), respectively. After 24 hours of pellicle biofilm development or sporulation, cells in the pellicle biofilms or sporulating cultures were harvested, diluted, and inoculated into fresh MSgg and DSM media again for a new round of competition. This process was repeated for four rounds. Total genomic DNAs from the co-cultures were also prepared after each round of competition. Using the genomic DNAs as the template, the *sda* gene was amplified by PCR, and the PCR products were analyzed with deep sequencing. This allowed us to identity the relative ratio of each engineered strain in the co-culture after each round based on the reads of the *sda* alleles.

For the biofilm competition assay (Figure 5A), a significant increase in the ratio of the *sda*^AGC^ allele in the co-culture was seen in both rounds 3 and 4. In contrast, the *sda*^TCC^ allele largely disappeared after round 1 (but detectable by sequencing). In the sporulation competition assay, significant changes in the ratio of the *sda*^AGC^ allele was seen even in round 1, and persisted throughout all 4 rounds, indicating that the strain bearing the *sda*^AGC^ allele quickly gained dominance in the sporulation media. The *sda*^AGT^ and *sda*^TCA^ alleles ranked equally as the second. Similar to what was seen in the biofilm competition assay, the *sda*^TCC^ allele disappeared quickly even from round 1 (Figure 5B). The above results suggest that in both biofilm and sporulation competition assays, the strain with the *sda*^AGC^ allele seemed to benefit from prolonged vegetative growth and thus outgrew other strains with different *sda* alleles. The prolonged vegetative growth in the *sda*^AGC^ strain is likely due to delayed Spo0A activation since Sda acts a checkpoint protein and governs the transition from vegetative growth to cell differentiation.

**Figure 5.**
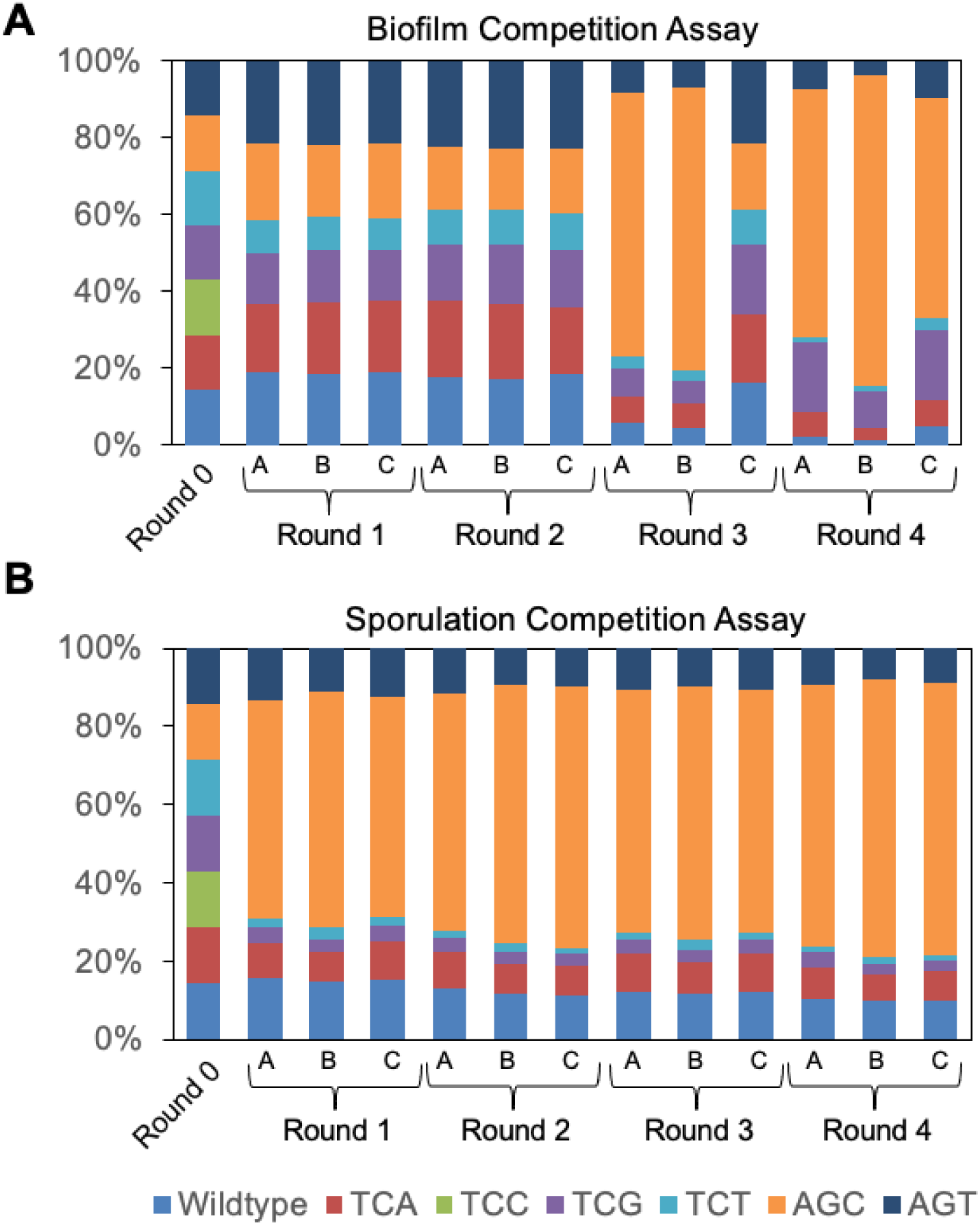
The *sda*^AGC^ allele outcompeted other alleles and the wild type in both biofilm and sporulation competition assays. **(A)** Results of the biofilm competition assays among the 7 complementation strains all with the *sda* gene deletion and complemented at *amyE* with either the wild type *sda* or one of the six synonymously substituted *sda* alleles. Cells of each strain were normalized for optical density and mixed together equally. The mixed culture was then diluted 1000-fold and incubated in MSgg medium at 30°C for 24 hours for pellicle biofilm development. Pellicle biofilm was collected and mildly sonicated to separate chains. Sonicated pellicle was then 1:1000 inoculated into 4 mL of fresh MSgg for another round of competition. This competition assay was repeated for a total of 4 rounds. In each round, total DNAs from the mixed culture were prepared and sent for deep sequencing to characterize the ratio of each *sda* alleles in the mixed culture. **(B)** Results of the sporulation competition assays among the 7 complementation strains described above. Cells of each strain were normalized for optical density and mixed together equally. Four rounds of sporulation competition assays were performed. In each round, the mixed culture was washed and resuspend in DSM broth, After 24 hours of shaking growth at 37°C, the mixed culture was heat-treated at 90°C for 20 min to kill vegetative cells. After cooling down at room temperature, the mixed culture was diluted 100-fold into fresh LB to allow spore germination. After that, the mixed culture was diluted 100-fold again to fresh DSM to start a new round of competition assay. After each round, total DNAs were prepared from the mixed culture and sent for deep sequencing. Assays were done in triplicate.

### Synonymously switching serine codons impacts protein levels of Sda

Synonymously switching serine codons in *sda* resulted in phenotypic changes in both biofilm formation and the timing of sporulation (at least in the case of AGC substitution). Based on our hypothesis, this could be due to altered Sda protein levels. In the previous study, we showed that systematically switching synonymous serine codons in *sinR* altered the SinR protein abundance accordingly, but not the *sinR* mRNA abundance in general (with a few exceptions)(Subramaniam *et al*., 2013).

We tested whether synonymously switching serine codons in *sda* also alters Sda protein abundance. We initially experienced some technical issues when using the Sda polyclonal antibodies to detect Sda by western blot (due to lack of specificity, data not shown). To bypass this issue, we fused each of the six *sda* alleles and the wild type *sda* to a *gfp* gene. Fusion proteins (Sda-GFP) expressed from the engineered *B. subtilis* strains can now be detected by commercially available polyclonal antibodies against GFP as well as by visualizing fluorescence of the cells under fluorescence microscopy. We picked three engineered strains expressing Sda^WT^-GFP, Sda^AGC^-GFP, and Sda^TCC^-GFP, respectively. Cells were grown in MSgg to early stationary phase (OD_600_=2.0) and fluorescence of the cells was compared. We also quantified fluorescence density of hundreds of individual cells in each of the three cell populations using MicrobeJ (Ducret *et al*., 2016). As shown in Figure 6A-B, cells expressing Sda^AGC^-GFP demonstrated the highest average pixel density while cells expressing Sda^TCC^-GFP had the lowest average pixel density.

**Figure 6.**
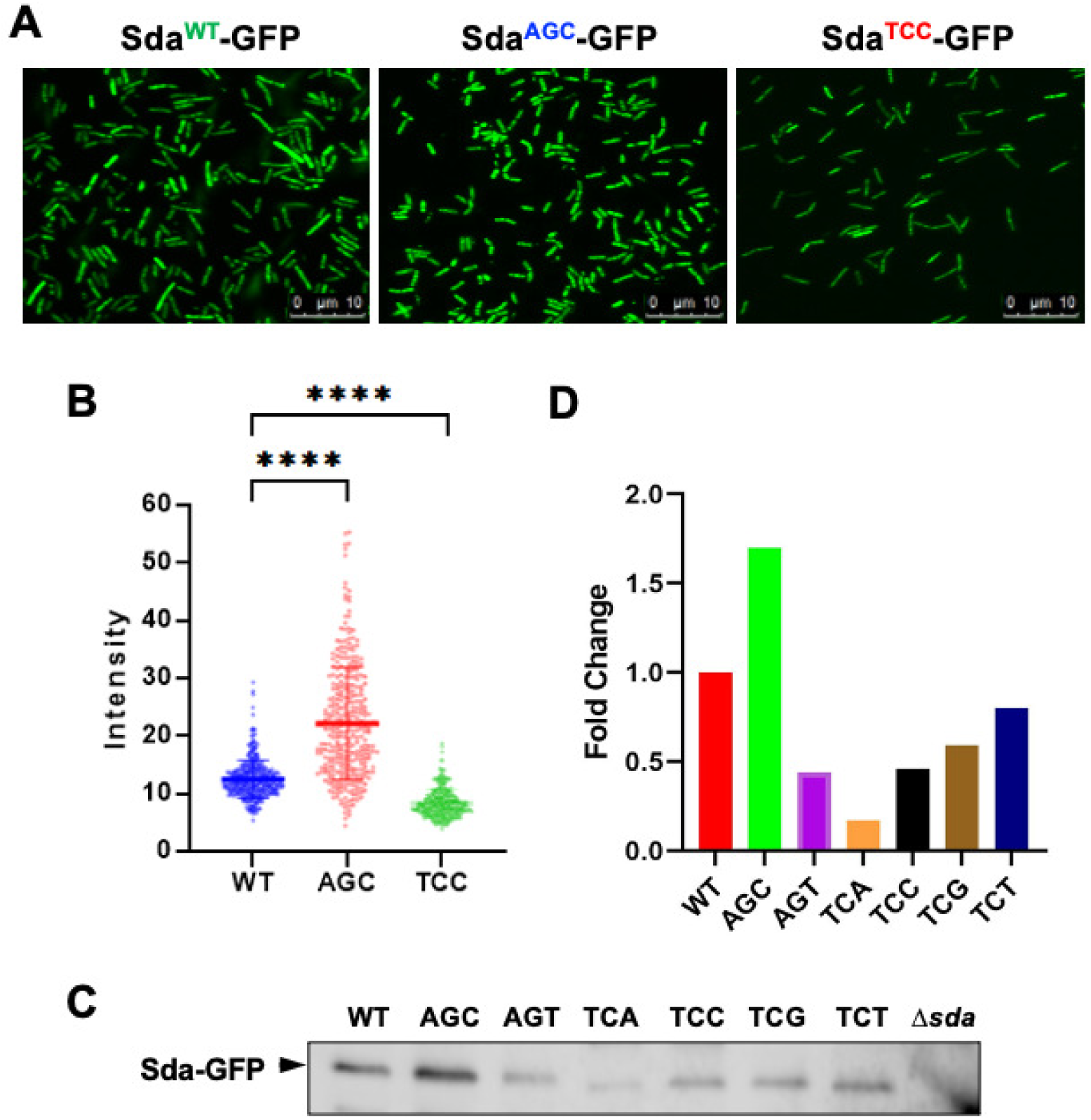
The *sda*^AGC^-*gfp* allele was expressed more abundantly than other serine codon synonymous alleles. **(A)** Cells of wild-type strain and strains with the *sda* serine codon synonymous alleles expressing Sda-GFP fusion proteins. Strains used here include the wild type (WT), and Δ*sda* complemented at *amyE* with either *sda*^AGC^-*gfp* or *sda*^TCC^-*gfp*. Fluorescent microscopic images were taken when cells reached stationary phase in MSgg medium. Scale bars shown are 10 μm. **(B)** Quantification of pixel density was conducted on about 300 individual cells from each of the 3 cell populations in (A) by MicrobeJ (Ref). **** indicates *p* value <0.0001. Statistics were performed by t-test. **(C-D)** Analyses of abundance of Sda-GFP fusion proteins by western blot. The 7 complementation strains all with the *sda* gene deletion and complemented at *amyE* with either the wild type *sda* or one of the six synonymously substituted *sda* alleles were used here. The *sda* strain was used as a negative control. Total protein lysates were prepared, normalized against the total protein concentration, and applied to SDS-PAGE. The Sda-GFP fusion protein was detected by polyclonal antibodies against GFP in western blot (C). Quantification of the pixel density of the protein bands from (C) was analyzed using the Image Lab software (Bio-Rad)(D).

We also preformed western blot to compare the abundance of the fusion proteins. As shown in Figure 6C-D, the level of Sda^AGC^-GFP (AGC) was about 1.7-fold higher than that of Sda^WT^-GFP (WT), whereas the rest of the fusion proteins all decreased to some degree compared to the wild type, with the most significance decrease seen in the Sda^TCA^-GFP fusion protein (TCA, 17% of the level of WT). To confirm that synonymously switching serine codons influenced translation but not transcription, we performed real-time quantitative PCR to measure the transcriptional levels of the fusion constructs in different strains when grown to either OD_600_=1 or 3 (log vs stationary phase). Our results showed no significant variations (Figure S3). In fact, the transcript for the *sda*^AGC^*-gfp* fusion even showed a lower abundance compared to the wild type (OD_600_=1, Figure S3), suggesting that elevated levels of the Sda^AGC^-GFP fusion protein (seen in Figure 6) were not likely due to transcription.

Taken all, our results supported the idea that increased Sda activities by synonymously switching the selected serine codons to AGC could be due to altered translation efficiency based on the serine codon hierarchy.

## Discussion

By applying global ribosome profiling, we previously found that under serine starvation, ribosomes pause frequently when translating the four UCN serine codons but not the two AGY codons on the mRNAs. We proposed a serine codon hierarchy (TCN vs AGC/AGT) based on the differential impacts that we observed, of the six synonymous serine codons on translation efficiency (Subramaniam *et al*., 2013, Greenwich *et al*., 2019). We hypothesized that this serine codon hierarchy could influence translation efficiency and thus activities of genes globally, and more importantly, differentially, depending on the ratio and bias toward the TCN serine codons in the genes. In addition, this global regulation is triggered by serine depletion as we previously showed (Subramaniam *et al*., 2013), suggesting that it acts as a regulatory mechanism in response to environmental changes. How can we determine if any gene in the *B. subtilis* genome (other than *sinR*) is subject to the regulation by the serine codon hierarchy? Genome-wide ribosome profiling could provide translation dynamics at the nucleotide-level resolution (Ingolia *et al*., 2012), but quite costly, and may still need validation by another method. In this study, by running a computer algorithm based on the proposed serine codon hierarchy, we generated a short list of 50 candidate genes that may be subject to this regulation. Furthermore, this bioinformatic approach identified *sinR* as enriched for the bad codons, which is already known to be subject serine codon usage-based regulation (Table 1 and Table S1).

Control of biofilm formation in *B. subtilis* is mediated by multiple regulators, including SinR, the key biofilm repressor, and other regulators that play either negative or positive roles (Figure 1). We demonstrated in the previous study that the serine codon hierarchy could function in a concerted manner during biofilm activation; genes for negative regulators such as SinR are at one end of the spectrum whose translation efficiency will be reduced upon serine depletion (due to higher ratios of the TCN serine codons) and therefore derepression of genes regulated by those negative regulators, whereas genes for positive regulators like Spo0A are at the opposite end of the spectrum (due to higher ratios of the AGY serine codons) that show sustained translation efficiency, and therefore, persistent activation of genes being regulated (Subramaniam *et al*., 2013). In this study, we provided evidence that another negative regulator in the same pathway, Sda, behaves similar to that of SinR (Figure 1).

Therefore, the two negative regulators SinR and Sda and the positive regulator Spo0A could act in a highly concerted fashion to ensure activation of biofilm matrix genes during biofilm induction. The observed impact of switching synonymous serine codons in genes like *sinR* and *sda* is indeed a manifest of what would have happened if the native genes have no bias in serine codon usage and translation of the mRNA does not respond to serine limitation.

Among the 50 candidate genes, many of them encode (known or predicted) secreted or membrane proteins (Table S1). We doubt this is just a pure coincidence. It is known that bacteria tend to use low-cost (or less costly) amino acids for the synthesis of secreted proteins since bacteria are less likely to recycle those proteins and reuse the amino acids (Smith & Chapman, 2010). Evidently, this type of selection (for low-cost amino acids) is a result of fitness during evolution. In our case, we don’t exactly know why those genes for secreted or membrane proteins contain a much higher ratio of TCN serine codons. One possibility we could think of is that this could serve as an energy saving strategy during serine depletion (a proxy for nutrient limitation) by more significantly reducing the synthesis of secreted or membrane proteins that are less likely or more difficult to be recycled by the bacteria.

## Methods and Materials

### Strains and media

Strains and plasmids used in this study are listed in Table 2. For general purposes, *Bacillus subtilis* strain PY79, 168, NCIB3610, and their derivatives were cultured in Luria-Bertani (LB) medium (10 g tryptone, 5 g yeast extract, and 5 g NaCl per liter broth) at 37°C. Concentrations of antibiotics that were added to media for *B. subtilis* strains were 10 μg ml^−1^ of tetracycline, 1 μg ml^−1^ of erythromycin, 100 μg ml^−1^ of spectinomycin, 20 μg ml^−1^ of kanamycin, and 5 μg ml^−1^ of chloramphenicol for transformation in *B. subtilis* and 100 μg ml^−1^ of ampicillin for *E. coli* DH5α transformations.

**Table 2.**
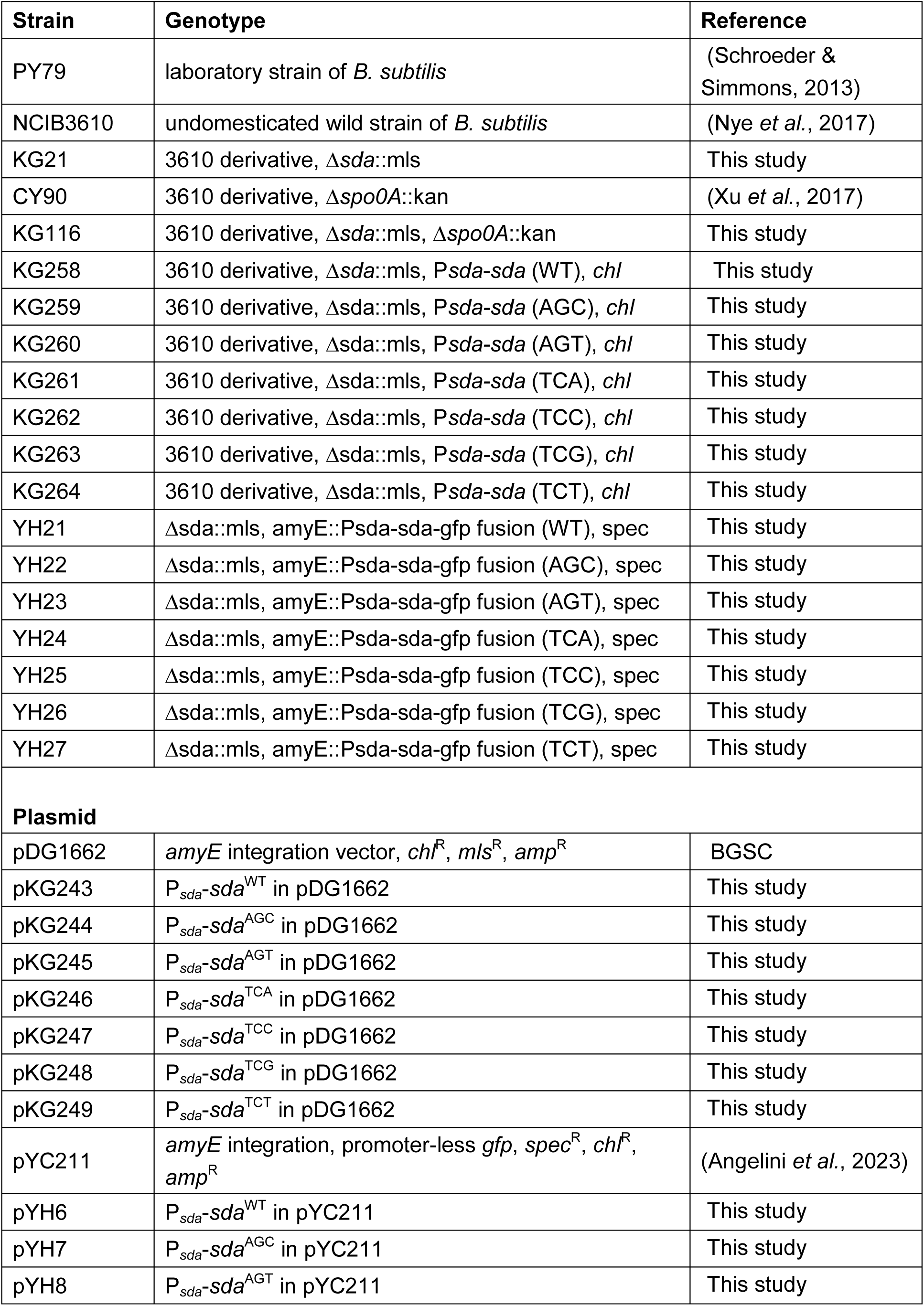
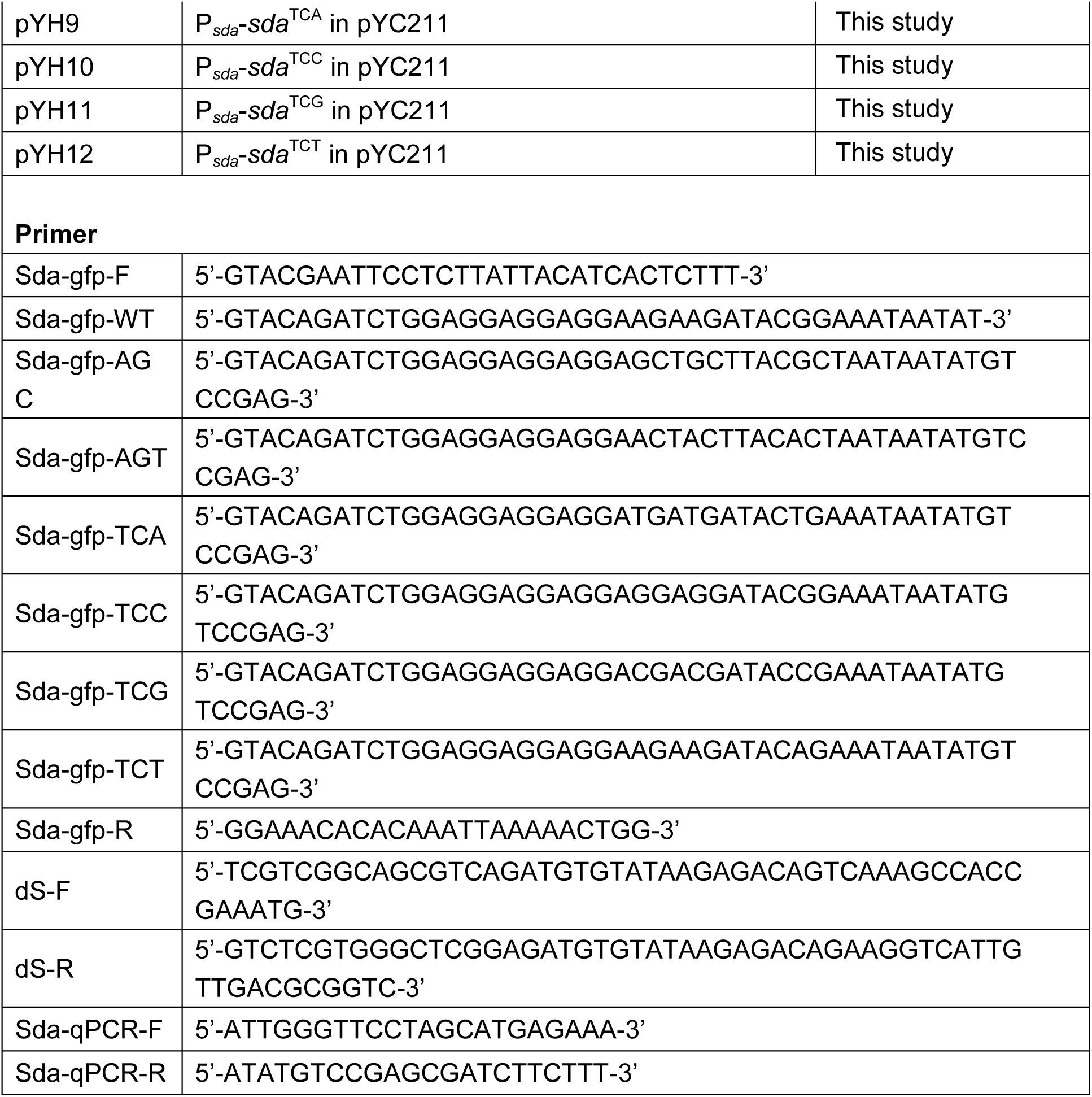
Strains, plasmids, and primers used in this study.

### Strain construction

Strain construction followed previously published protocols for molecular cloning and transformation (Qin *et al*., 2019). The *sda* gene deletion strain in 168 background was purchased from the *Bacillus* Genetic Stock Center (BGSC, http://www.bgsc.org) and the gene deletion was then introduced NCIB3610 by transformation. The wild-type complementation strain was constructed by transforming the plasmid pDG1662 cloned with the wild-type *sda* gene and integrate P*_sda_*-*sda*^WT^ into the *amyE* locus of the Δ*sda* strain. To construct synonymous serine mutants, the *sda* alleles with indicated synonymous mutations at the terminal three serine codons were similarly constructed in the plasmid pDG1662 and introduced into the *amyE* locus of the Δ*sda* strain. The *sda-gfp* gene fusion (wt *sda* or *sda* with indicated synonymous mutations) were generated using the same method, but by using the plasmid pYC211 (Angelini *et al*., 2023). Other strains were constructed by applying synthetic DNA fragment or site-directed mutagenesis. The *sda-gfp* translational fusions (wild-type *sda* or *sda* with indicated synonymous substitutions in the terminal three serine codons) were amplified using primers sda-gfp-F and sda-gfp-R and using DNAs containing wild type or *sda* alleles as templates. The PCR products were then digested using *Bgl*II and *EcoR*I, purified by gel electrophoresis, and ligated into the plasmid pYC211 treated with the same digestions.Ligations were transformed into *E. coli* and recombinant plasmids were prepared and verified by DNA sequencing. Recombinant plasmids were then introduced into *B. subtilis sda* mutant by transformation. Transformants were selected on LB agar plates supplemented with appropriate antibiotics and integration of *sda-gfp* at *amyE* was verified by sequencing.

### Bioinformatic approach to identify high-TCN usage genes

The genome of *Bacillus subtilis* NCIB 3610 (accession number NZCM000488.1) was accessed from NCBI. For each protein-coding gene in the genome, each of the 64 codons was counted through the gene. To determine which genes were coded with the “bad” serine codons (TCN), a scaling factor was multiplied to the raw counts of each serine codon in the gene. To root the hierarchy in a quantifiable ranking, the fold-enrichment of ribosome density observed during serine starvation conditions at each of the six serine codons was used (see Fig. 3C in the reference)(Subramaniam *et al*., 2013). For TCT, TCC, TCA, TCG, AGT and AGC codons, the scaling factor was 1.31, 1.78, 1.24, 1.87, 0.88, and 0.75 respectively, which gives a higher ranking to the TCNs over the AGY codons.

The raw counts of each codon were by the scaling factor and the score was totaled for each gene. This raw score was then normalized by the length of the gene, resulting in an adjusted, normalized score of serine codon usage. All genes less than 50 codons in length were considered small genes and removed from the final top 50 ranking.

### Biofilm assay

Cells were grown in LB broth at 37°C for 4 hours, and then normalized to OD_600_=1.0. Three μL of the culture was inoculated into 3mL of MSgg liquid media in a 12-well microtiter plate. All plates for biofilm assays were incubated at 30°C for 1-2 days to allow pellicle development. Images of pellicle biofilms were taken by using a Leica MSV269 dissecting microscope with a Leica DFC2900 camera and x4 magnification under the same exposure and acquisition settings.

### Membrane staining

Overnight cultures grown in LB broth at 37°C were normalized to OD_600_=1.0. Overnight cultures were 1:100 diluted into 3 mL DSM media and grown to OD_600_=1.0 again. Next, in a 50mL Erlenmeyer flask, 200 μL of the DSM culture was inoculated into 20 mL of DSM liquid media and incubated at 37°C. These steps were to improve the synchronization of sporulation of the cells. After 6, 7, and 8 hours of incubation, respectively, 500 μL of the culture was transferred into a 1.5 mL micro tube and then centrifuged at 5000 rpm for 1 min. Supernatant was discarded, and the cell pellet was suspended with 50 μL of sterile water. One μL of the FM4-64 dye (Thermo-Fisher) was mixed with 50 μL of the suspended culture, and the micro tube was wrapped with aluminum foil and placed on ice for 1 min. Cells were then imaged by using a Leica DFC3000 G camera on a Leica AF6000 microscope. Samples from all time points were taken under the same exposure and image acquisition settings.

### Sporulation assay

Fresh cultures of wildtype, AGC, and TCC strains were grown in LB shaking at 37°C until reaching OD_600_=1. Then, the culture was 1 to 100 inoculated into a flask containing 15ml DSM medium and incubated in the shaker at 37°C. Partial cell culture was collected at 6, 8, and 10 hours of incubation respectively to determine the sporulation rate. Collected culture, which contained live cells and spore, was serial diluted and plated on LB agar plate before heat-killing. Then, the remaining collected cultures were incubated in 80°C water bath for 25 minutes and cooled in ice for 10 minutes to kill all live cells. Viable spores remained in the culture was serial diluted and plated on LB agar plate as well. Cells on the plates were quantified after overnight incubation at 37°C. Terminal sporulation rate of the wild-type strain and all synonymous serine mutants were obtained following the protocol described above after 24 hours of sporulation. The heat-kill assay was carried out following the published protocol (Angelini *et al*., 2023).

### Western blot

Twenty mL cultures were grown at 37°C and harvested when the cultures reached OD_600_=2.5. Cells were spun down at 4000 rpm for 10 min. After discarding supernatant, 10 mL of lysis buffer (20 mM Tris-HCl, 200 mM NaCl, 1 mM EDTA pH 7.4) was used to wash the cell pellets once. Cell pellets were resuspended in 1.2 mL lysis buffer and incubated with 20 μg/mL of lysozyme (NEB) for 30 min on ice. Samples were sonicated on ice 3 times for 30 sec each. Sonicated and lysed cells were centrifuged (14000 rpm, 30 min, 4°C) to remove cell debris. Cleared lysates were normalized for total protein concentrations using Bradford assay kit (Thermo-Fisher, Waltham, MA), and were run on an SDS-PAGE that consisted of 15% separating gel (2.4 mL H_2_O, 5 mL acrylamide (30 %), 2.5 mL 1.5 M tris (pH 8.8), 50 µL SDS (20%), 100 µL APS (10%), 10 µL TEMED) and 6% stacking gel (2.7 mL H_2_O, 0.8 mL acrylamide (30 %), 0.5 mL 1 M Tris (pH 6.8), 20 µL SDS (20%), 40 µL APS (10%), 4 µL TEMED). Once electrophoresis was complete, proteins on the SDS-PAGE were transferred to a PVDF membrane (Millipore, Billerica, MA) at 100V for 3 hours at 4°C. The membrane was washed once with TBS buffer, and then blocked with 20 mL TBS buffer mixed with 5% skin milk for an hour. Then, the membrane was incubated with the polyclonal anti-GFP antibody (1:5000, Abcam) overnight at 4°C. After washing with TBS buffer 3 times for 5 min each, the membrane was incubated with the goat-anti-rabbit secondary antibody (1:10,000, Bio-Rad). After incubation, the membrane was washed with TBS buffer 3 times for 5 min each. The membrane was incubated with SuperSignal West Dura chemiluminescent substrate (Thermo-Fisher, Waltham, MA) for 15 min in dark and was imaged on a ChemiDoc Imaging System (Bio-Rad). Intensities of the resulting bands on the membrane were detected and analyzed by using Image Lab™ Software (Bio-Rad).

### Quantitative real-time PCR

Cultures were inoculated in 20 mL of DSM liquid media. Once cultures were grown to exponential phase (OD_600_=1.0) or stationary phase (OD_600_=2.5), cells were harvested and spun down (15000 rpm for 5 min). One mL of TRIzol reagent (Thermo-Fisher) was added to resuspend the cell pellets, and the resuspended cells were incubated for 5 min at room temperature. Total RNAs were extracted by using the Zymo Direct-zol RNA kit (Invitrogen) and following the protocol provided by the manufacturer.

Concentrations of the extracted RNAs were measured by using a NanoDrop One instrument (Thermo-Fisher). The extracted RNAs were then reversely transcribed into cDNA by using a high-capacity cDNA reverse transcription kit (Applied Biosystems). Primers for quantitative PCR were listed in Table 2. RT-qPCR was run by using Fast SYBRTM Green Master Mix (Applied Biosystems) and a Step One Plus Real-Time PCR system (Applied Biosystems). The *sigA* gene served as an internal reference for calculating relative expression of genes of interest using the 2^−ΔΔCT^ method (Qin *et al*., 2019).

### Sporulation competition assay

The sporulation competition assay (SCA) was performed by co-culturing in DSM the 7 complemented strains with either the WT *sda* gene and or each of the six synonymous *sda* alleles. Cultures were grown to OD_600_=1.0, and 20 μL of each strain were mixed and added to 4 mL of DSM liquid media (200x dilution). The mixed culture was grown at 37°C for 24 hours and heat-treated at 90°C for 20 min to kill all vegetative cells. After cooling down at room temperature, the heat-treated culture was 100-fold diluted. 20 μL of the diluted culture was transferred into 4 mL of LB broth at 37°C for 4 hours so that spores in the mixed culture were germinated. Twenty μL of the regrown culture was inoculated into 4 mL of fresh DSM for the next round of competition. A total of four rounds of competition were performed.

### Biofilm competition assay

The biofilm competition assay (BCA) was performed similarly from above. Each of the 7 complemented strains were grown in LB to OD_600_=1.0. Twenty μL of each of the 7 cultures was mixed together, added into 6 mL of MSgg media, and incubated at 30°C for 24 hours to allow pellicle biofilm development. The pellicle biofilm was then transferred to 4 mL of PBS buffer. Chains in the pellicle were first sheared by pipetting and then mildly sonicated on ice 3 times for 15 sec each. Sonicated culture was diluted 100-fold, and 60 μL of the diluted culture was inoculated into 6 mL of fresh MSgg liquid media to initiate second round of pellicle development and competition assay. Two mL of the regrown culture and the sonicated culture from each round of the competition assay were harvested for DNA extraction and the DNAs were applied for deep sequencing. In addition, both the regrown culture from SCA and the sonicated cultured from BCA were applied for colony-forming unit (CFU) assay. For each round of competition assay, 10 colonies were selected for Sanger sequencing on *sda* as another control.

### DNA sequencing

The *sda* gene of the mixed strains sample from sporulation and biofilm competition assay were amplified by using two primers (dS-F:5’-TCGTCGGCAGCGTCAGATG-TGTATAAGAGACAGTCAAAGCCACCGAAATG-3’ and ds-R:5’-GTCTCGTGGGCT-CGGAGATGTGTATAAGAGACAGAAGGTCATTGTTGACGCGGTC-3’). Thermal cycling of PCR was run using Phusion High-Fidelity PCR Master Mix with HF Buffer (ThermoFisher, USA) for 30 cycles. The PCR products were purified using GeneJET PCR Purification Kit (ThermoFisher, USA), and then applied for sequencing library construction by using the KAPA library preparation kit (Kapa, MA, USA) and following the manufacturer’s instructions. Finally, the library was analyzed for quality by the Agilent Bioanalyzer 2100 system and then sequenced on the HiSeq2500 platform (Illumina, CA, USA).

### Data analysis

FLASH (Fast Length Adjustment of Short reads), an analysis tool that merge paired end reads, was used to stitch the paired end reads from the sequenced amplicons if the reads align with each other. Meanwhile, reads that fail to assemble were discarded. The amount of each strain was recognized and quantified according to their unique serine sequence.

## Acknowledgments

We are grateful to Dr. Alan Grossman (MIT) for the gift of polyclonal antibodies for Sda and *sda* strains used in our initial tests. We would also like to thank members in the Chai and Godoy laboratories for helpful discussions and suggestions during this study. This work was supported by an NSF grant to Chai Y (MCB 1671532), and a grant from National Natural Science Foundation of China to Qin Y (32102381).

## Author contributions

YH, YQ, KG, and YC designed the study. YH, YQ, JG, SB, MD, and KG performed the experiments. KG designed algorism and performed codon analyses. YH, YQ, KG, and YC did data analyses. YH, KG, and YC wrote the manuscript.

**The authors declared no conflict of interest.**

